# Physiological motion of the carotid atherosclerotic plaque quantified using ultrasound B-Mode image analysis

**DOI:** 10.1101/255232

**Authors:** Baris Kanber, Timothy C. Hartshorne, James W. Garrard, A. Ross Naylor, Thompson G. Robinson, Kumar V. Ramnarine

## Abstract

**Background:** Physical motion throughout the cardiac cycle may contribute to the rupture of the atherosclerotic carotid plaque, resulting in ischaemic stroke. The purpose of this study was to quantify the physiological motion of the atherosclerotic carotid plaque and to investigate any relationship between the quantified motion parameters and the degree of stenosis, greyscale plaque characteristics, and the presence of cerebrovascular symptoms.

**Methods:** Displacement, velocity and acceleration of 81 plaques (51% symptomatic, stenosis range 10%-95%) from 51 patients were measured using an automated system employing a block matching algorithm relative to the ultrasound probe and relative to the periadventitial tissues, over a mean duration of 5 cardiac cycles.

**Results:** Averaged across all plaques, the displacement amplitude was 1.2 mm relative to the probe, and 0.35 mm relative to the periadventitial tissues. Maximum and mean plaque velocities were 4.7 and 1.3 mm/s relative to the ultrasound probe, and 2.4 and 0.70 mm/s relative to the periadventitial tissues. The corresponding acceleration magnitudes were 69 and 22 mm/s^2^ relative to the probe, and 57 and 18 mm/s^2^ relative to the periadventitial tissues. There were no significant differences in any of the motion parameters, with respect to the presence of cerebrovascular symptoms, and none of the parameters showed a statistically significant relationship to the degree of stenosis, and the greyscale plaque characteristics (p≥0.05). The technique used was able to detect plaque motion amplitudes above 50μm.

**Conclusions:** This study provides quantitative data on the physiological motion of the atherosclerotic carotid plaque *in-vivo*. No significant relationship was found between the measured motion parameters and the presence of cerebrovascular symptoms, the degree of stenosis, and the greyscale plaque characteristics.

## Introduction

Ultrasound imaging is routinely used to assess atherosclerotic plaques in the carotid arteries. Although the processes increasing plaque vulnerability are not fully understood, it is thought that physical motion may play a part, contributing to plaque rupture and embolisation [1-5]. Studies investigating plaque motion using B-Mode ultrasound have mainly employed qualitative methods [6-7], except for scarce conference proceedings [8-9] and studies utilising radiofrequency data [5,9]. Iannuzzi et al., for example, quantitatively assessed longitudinal lesion motion and found an association with ipsilateral brain involvement in transient ischaemic attack (TIA) patients. Other investigators found relationships between recurrent ischaemic attacks and the motion of the intra-plaque contents [1,10]. Meairs and Hennerici measured maximal plaque velocities using three dimensional ultrasound [11]. Other studies measured the motion of the carotid artery instead of the plaque [12-25], while some investigators assessed intra-plaque strains [26-33]. However, previous studies have not provided a comprehensive evaluation of plaque motion, including physical plaque accelerations, relative to the ultrasound probe and relative to the periadventitial tissues using equipment readily available in a typical stroke clinic [34]. Therefore, the purpose of this study was first to undertake an evaluation of carotid plaque motion relative to the ultrasound probe (bulk motion) and relative to the periadventitial tissues using a clinical scanner, and secondly, to investigate any relationships between the measured motion parameters and the degree of stenosis, greyscale plaque characteristics, and the presence of cerebrovascular symptoms. Our hypothesis was that quantitative assessment of carotid plaque motion using a clinical scanner can identify the “at-risk” carotid plaque at the bedside.

## Methods

Fifty-one patients (61% male, mean age 77 years) who attended the University Hospitals of Leicester NHS Trust's Rapid Access Transient Ischemic Attack (TIA) clinic were recruited for this study. The study was approved by the National Research Ethics Service (NRES) Committee East Midlands - Northampton (reference 11/EM/0249) and followed institutional guidelines. Patients gave written, informed consent before participating in the study. Ultrasound image sequences of the carotid plaque in the longitudinal plane were acquired by experienced sonographers using a Philips iU22 ultrasound scanner (Philips Healthcare, Eindhoven, The Netherlands) and an L9-3 probe. B-Mode (greyscale) image sequences were recorded as DICOM files over an average duration of 5.6 seconds (mean frame rate was 32 frames per second) using the vascular carotid preset on the scanner (Vasc Car preset, persistence low, XRES and SONOCT on). Motion analysis was carried out for 81 plaques (stenosis range 10-95%). Each plaque was classified as either having caused cerebrovascular symptoms in the ipsilateral brain hemisphere within the past six-month period (i.e. symptomatic) or as asymptomatic following independent, specialist medical review. The symptomatic plaque was used as a surrogate marker of the unstable carotid plaque.

Laboratory experiments to validate the motion analysis were carried out using an actuator device that generated programmable and repeatable periodic displacements of a tissue mimicking material (TMM). The specifications of this precision lead-screw-based linear actuator (T-LA-28S, Zaber Technologies Inc, Richmond, British Columbia, Canada) indicated a typical linear displacement accuracy of 8 μm and a precision of 0.3 μm The actuator movement was controlled by a desktop computer through the RS-232 interface and was programmed to produce a more rapid displacement of the posterior wall away from the probe (downward) compared to the displacement towards to probe (upward) in order to mimic the rapid dilation of the carotid artery during systole compared to the less rapid relaxation during diastole [23]. The actuator moved a Perspex plate cut-out measuring 4 centimetres in diameter that formed part of the bottom of a water tank. The bottom was covered with a watertight latex membrane sealed with silicone, which allowed vertical movement of the Perspex plate by the actuator. There was no programmed motion in the horizontal direction. A test object made of a rectangular block of TMM was placed on the Perspex plate and used for motion analysis (Figure 1). The TMM used was an agar-based formulation, which had good acoustic properties and met the requirements of the IEC 1685 draft report [35-36]. The composition (by weight) was 82.97% water, 11.21% glycerol, 0.46% benzalkoniumchloride, 0.53% SiC powder (400 grain, Logitec Ltd., Glasgow, UK), 0.94% Al2O3 powder (3 μm Logitec), 0.88% Al2O3 powder (0.3 μm Logitec), and 3.00% agar (Merck Eurolab). The TMM was prepared by heating the ingredients to 96°C (±3°C) for 1 hour using a double boiler whilst stirring continuously with a motorised stirrer. The mixture was then allowed to cool down to 42°C and cast into shape. DICOM image sequences were recorded with the actuator set to produce maximum displacements of 5, 10, 20, 50, 100, 200 and 500 μm.

**1 -.**
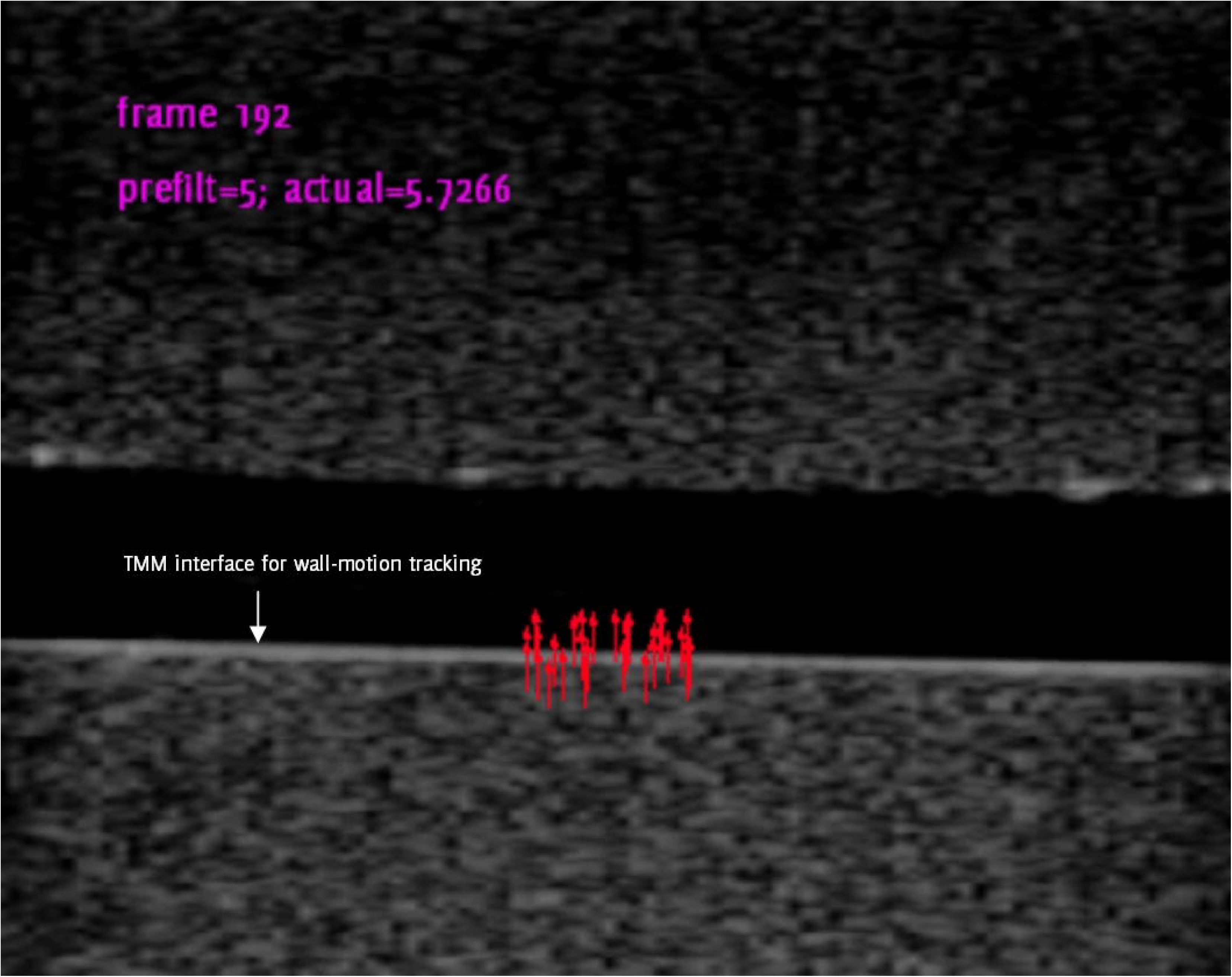
A still frame from an image sequence of the tissue mimicking material (TMM, below) with the actuator set to produce a maximum of 500 μm displacement from the initial position. Red arrows show the local TMM displacement at time t relative to the position at frame 1, magnified by a factor of 10. White arrow shows the TMM interface used for wall motion tracking (see Methods/Motion Tracking). The top block of TMM was attached to the ultrasound probe and was stationary.

Quantitative analyses were carried out using MATLAB version 7.14 (MathWorks,Natick, Massachusetts, USA) and SPSS version 20 (IBM Corporation, Armonk, New York, USA). The degree of carotid artery stenosis (DOS) was measured using criteria consistent with the NASCET method utilizing blood flow velocities in conjunction with the B-Mode and colour flow imaging [37-39]. Normalized plaque greyscale median (GSM) and surface irregularity index (SII) were measured using previously described methods [40-41] and were averaged over all the ultrasound image frames acquired for each artery.

### Motion Assessment

Motion assessment was performed using a block matching algorithm with the normalised correlation coefficient as the similarity measure [40]. Several points were manually selected in the first frame of the ultrasound image sequence at an approximate, uniform grid spacing of 1/2 mm covering the whole plaque body, or a 1 mm thick slab of tissue in the case of periadventitial tissues. These points were then automatically tracked in the successive image frames by iteratively testing template sizes from 20x20 mm^2^ down to 6x6 mm^2^ in decrements of 1x1 mm^2^ (Figure 2).

**2 -.**
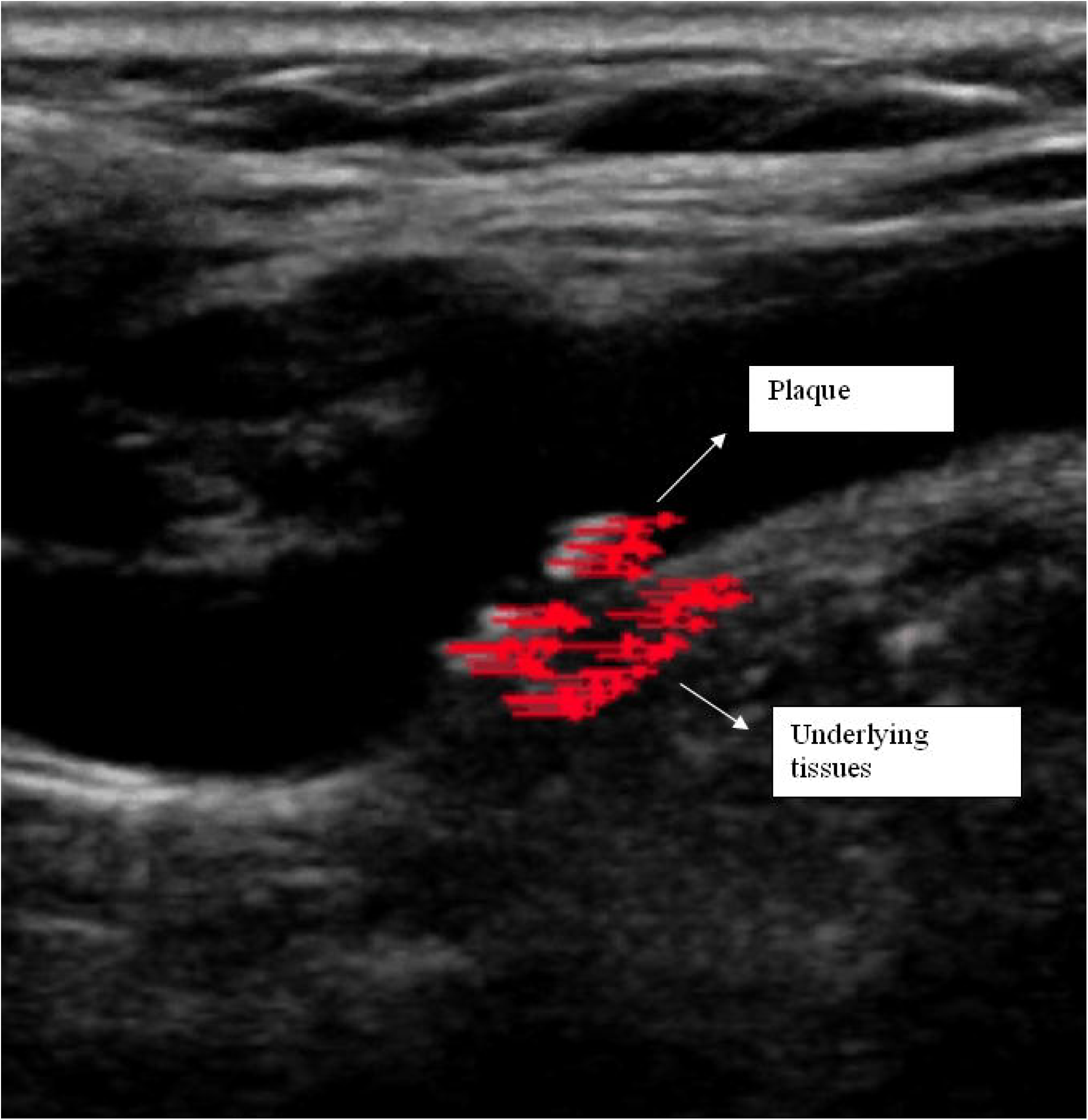
Plaque and periadventitial tissues with motion tracking. Arrows show the local displacement at time t with respect to the position at frame 1, magnified by a factor of 10. At this point in time, the plaque and the periadventitial tissues are seen to have moved uniformly in the horizontal direction.

The individual points describing the motion of the plaque and the periadventitial tissues were separately averaged, resulting in two motion trajectories; one for the plaque (*r*_plaque_) and one for the periadventitial tissues (*r*_tissue_). Motion of the plaque relative to the periadventitial tissues (*r*_rel_) was calculated by transforming *r*_plaque_ to a reference frame the origin of which moved according to *r*_tissue_. Maximum displacement was calculated as the furthest distance between any two points assessed over all possible pairs of points on a given trajectory, while plaque velocity and acceleration were determined as the first and second derivatives, respectively, of the motion trajectories using the central difference method (gradient function in MATLAB). Reproducibility was assessed by measuring each motion parameter 5 times for 10 plaques, and subsequently determining the intra-observer coefficients of variation. The same ultrasound image sequences were used for the reproducibility analysis; thus no additional scans were performed. In the case of the *in vitro* analysis, motion parameters obtained using the present technique were compared against those obtained using a previously described method, which tracked the boundary of the tissue mimicking material (Figure 1) [42-43].

### Statistical Methods

The non-parametric Wilcoxon-Mann-Whitney test was used to determine whether motion parameters differed significantly between plaques that were or were not associated with symptoms. Logistic regression testing, one for the motion parameters relative to the ultrasound probe, and one for the motion parameters relative to the periadventitial tissues, were further employed to assess whether any of the motion parameters were significant predictors of symptoms. Correlations between the motion parameters and the degree of stenosis, plaque greyscale median and the surface irregularity index were determined using Spearman's rank correlation coefficient. In all cases, two-tailed p-values less than 0.05 were considered statistically significant.

## Results

There were 31 male patients and 20 females, aged between 58 and 95 years. Prevalence of the cardiovascular risk factors were as follows: 67% hypertension, 55% hypercholesterolaemia, 27% ischaemic heart disease, 16% atrial fibrillation, 27% diabetes, 43% previous TIA/stroke, 8% peripheral vascular disease, 65% history of smoking, 37% alcohol consumption, and 29% family history of stroke.

Motion analysis was successful for 66 plaques (81%) and unsuccessful for 15 (19%). The causes of motion tracking failure were speckle decorrelation (13 plaques), and ultrasonic shadowing (2 plaques). The mean normalized correlation coefficient was 0.95 for the plaques for which motion tracking was successful. Figure 2 shows an example still frame (and Additional Video File, the corresponding video) for a plaque with successful motion tracking, while Figure 3 shows the calculated motion parameters for the same plaque, relative to the ultrasound probe. Cardiac cycles are clearly visible in the plots showing the horizontal and vertical components of plaque position. They are also visible, albeit less clearly, in the plots showing the plaque velocity and acceleration. The drift in the plot showing the horizontal component of plaque position is thought to be due to patient/bulk tissue motion (relative motion between the ultrasound probe and the patient).

**3 -.**
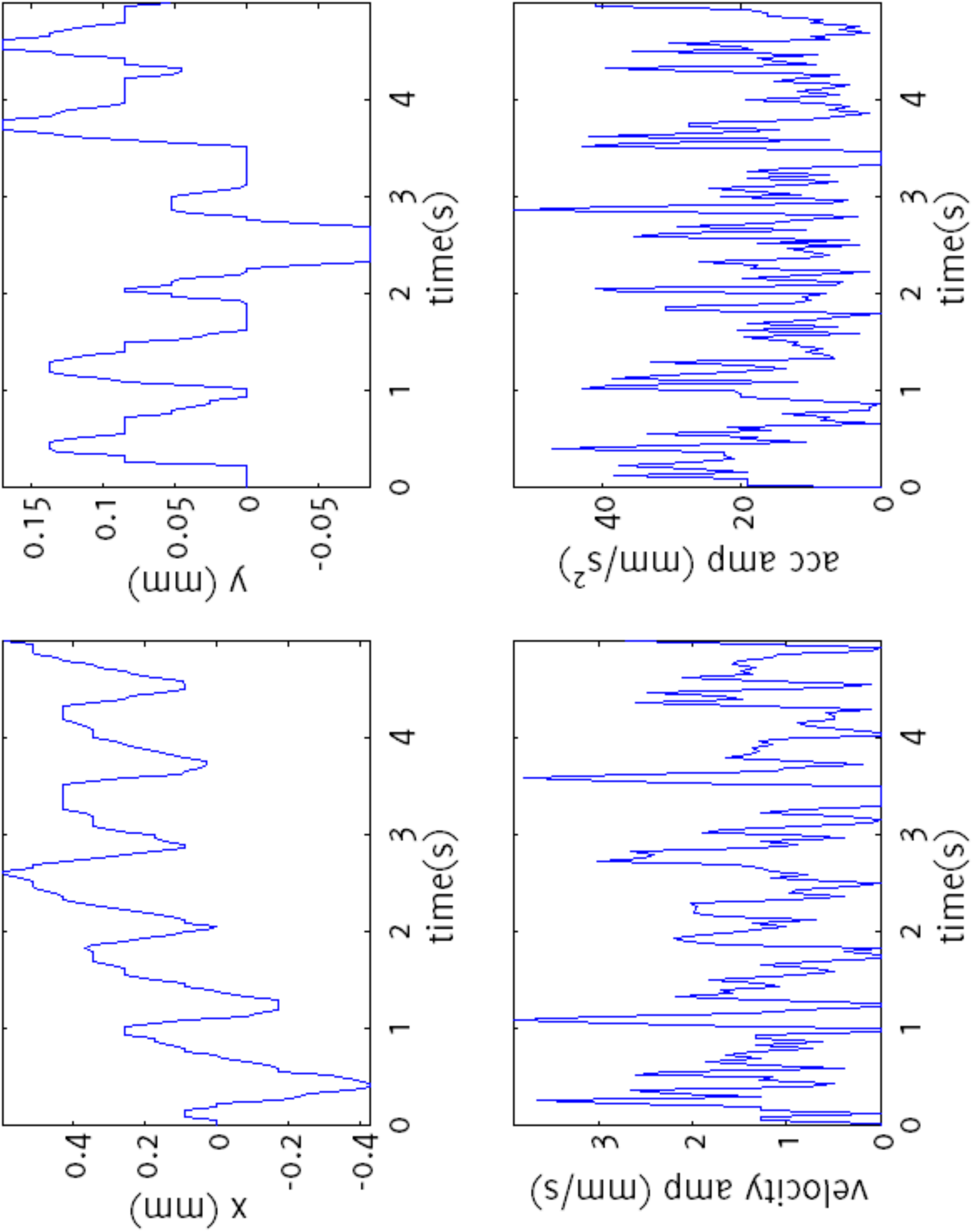
Calculated motion parameters, relative to the ultrasound probe, for the plaque sample shown in Figure 2. The horizontal and vertical components of plaque position are shown in the top-left and top-right plots, respectively. Velocity and acceleration magnitudes are shown in the plots at the bottom.

The mean values across all plaques of the motion parameters relative to the probe and the periadventitial tissues were as shown in Tables 1 and 2. Averaged across all plaques, the displacement amplitude was 1.2 mm relative to the ultrasound probe, and 0.35 mm relative to the periadventitial tissues. Maximum and mean plaque velocities were 4.7 and 1.3 mm/s relative to the ultrasound probe, and 2.4 and 0.70 mm/s relative to the periadventitial tissues. Maximum and mean plaque accelerations were 69 and 22 mm/s^2^ relative to the probe, and 57 and 18 mm/s^2^ relative to the periadventitial tissues. For the motion relative to the periadventitial tissues, average displacement amplitude was low (less than 0.4 mm) and the differences between the symptomatic and asymptomatic groups were not statistically significant (Table 2 and Figure 4). In the case of the motion of the plaque relative to the probe, the average displacement amplitude was much larger (>1 mm) but the differences between the symptomatic and asymptomatic groups were also statistically insignificant (Table 1 and Figure 4). Logistic regression testing further confirmed that none of the motion parameters, either relative to the ultrasound probe or relative to the periadventitial tissues, were significant predictors of the presence of cerebrovascular symptoms (p>0.05). There was no statistically significant association between any of the motion parameters and the degree of stenosis, or the greyscale plaque characteristics (Tables 3 and 4).

**1 -.**
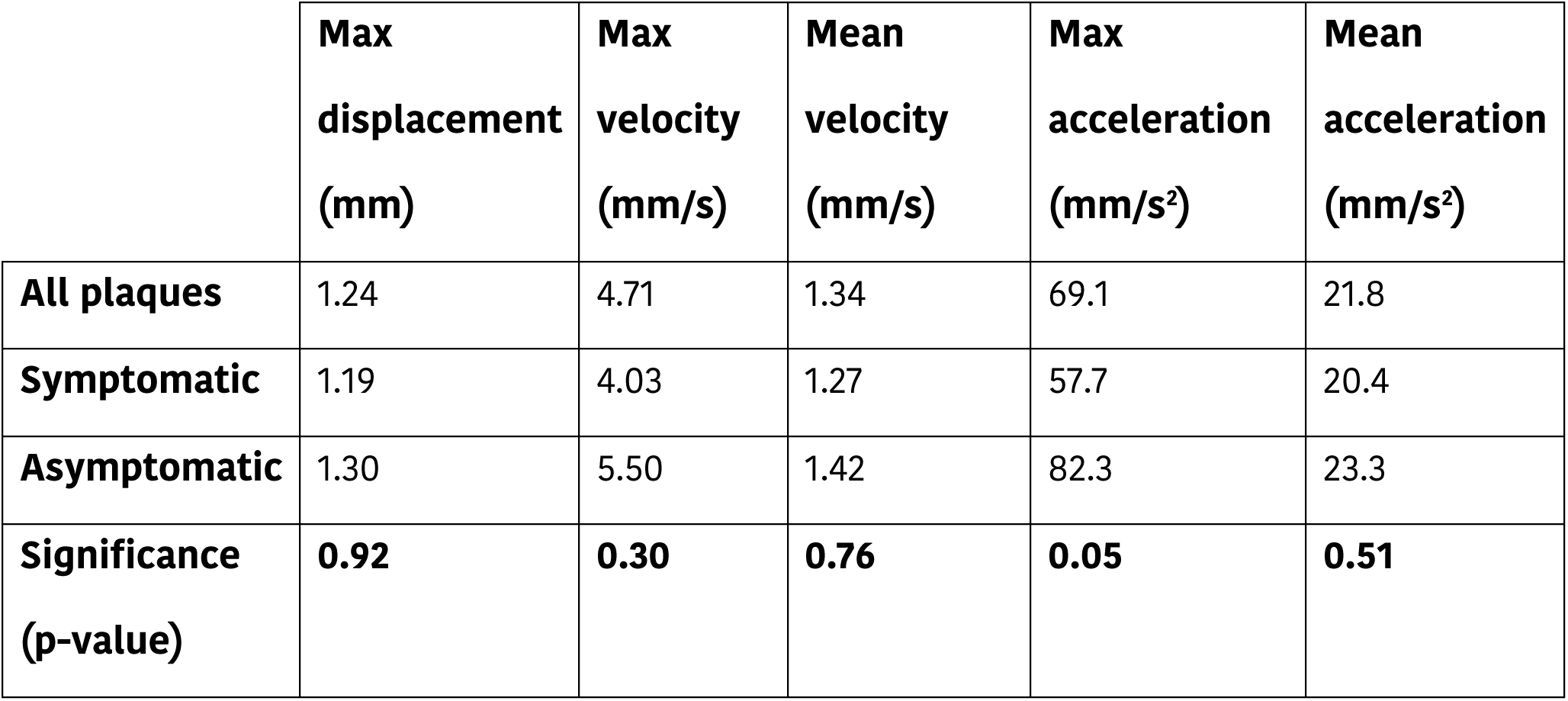
Mean values, across plaques, of the motion parameters relative to the ultrasound probe.

**2 -.**
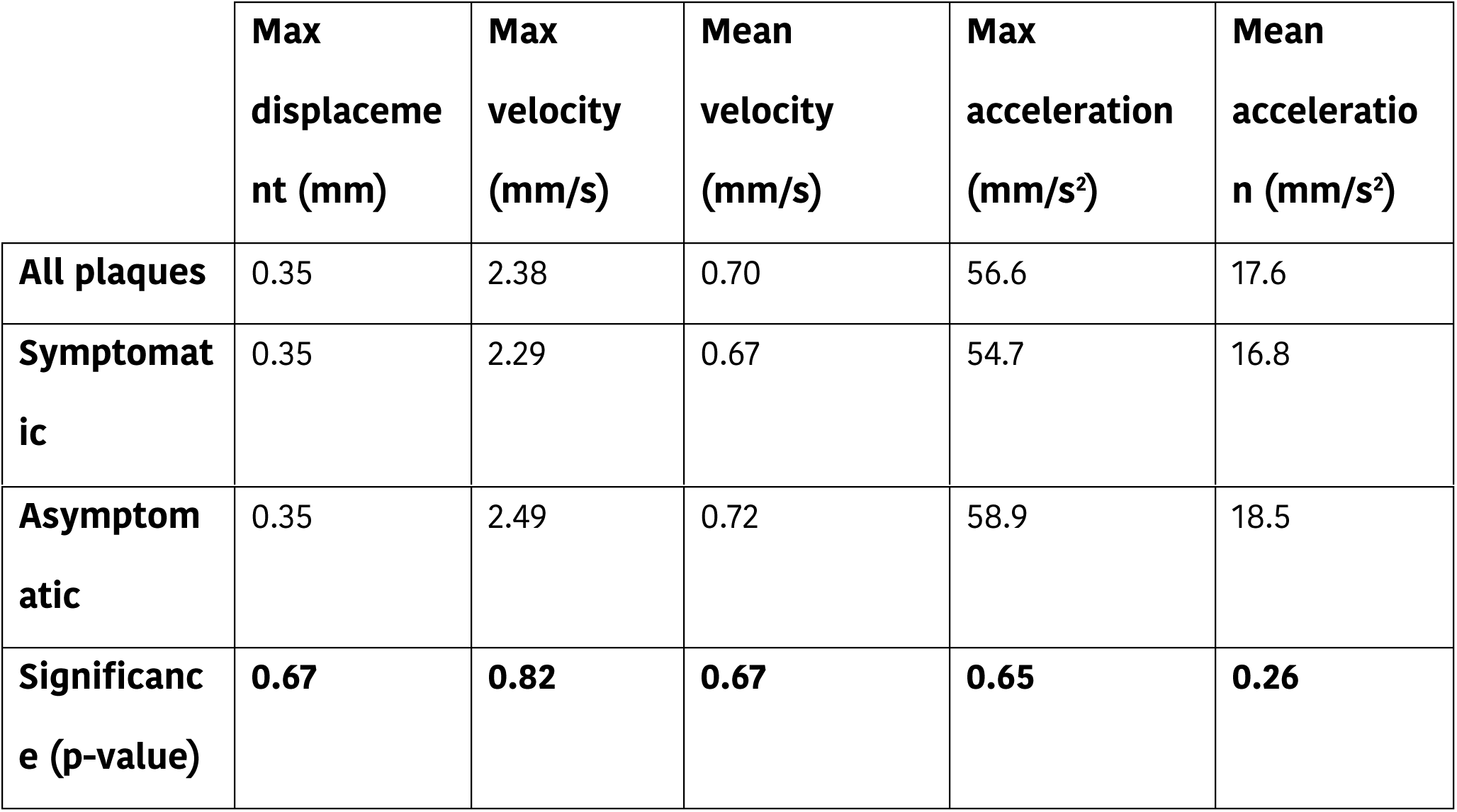
Mean values, across plaques, of the motion parameters relative to the periadventitial tissues.

**4 -.**
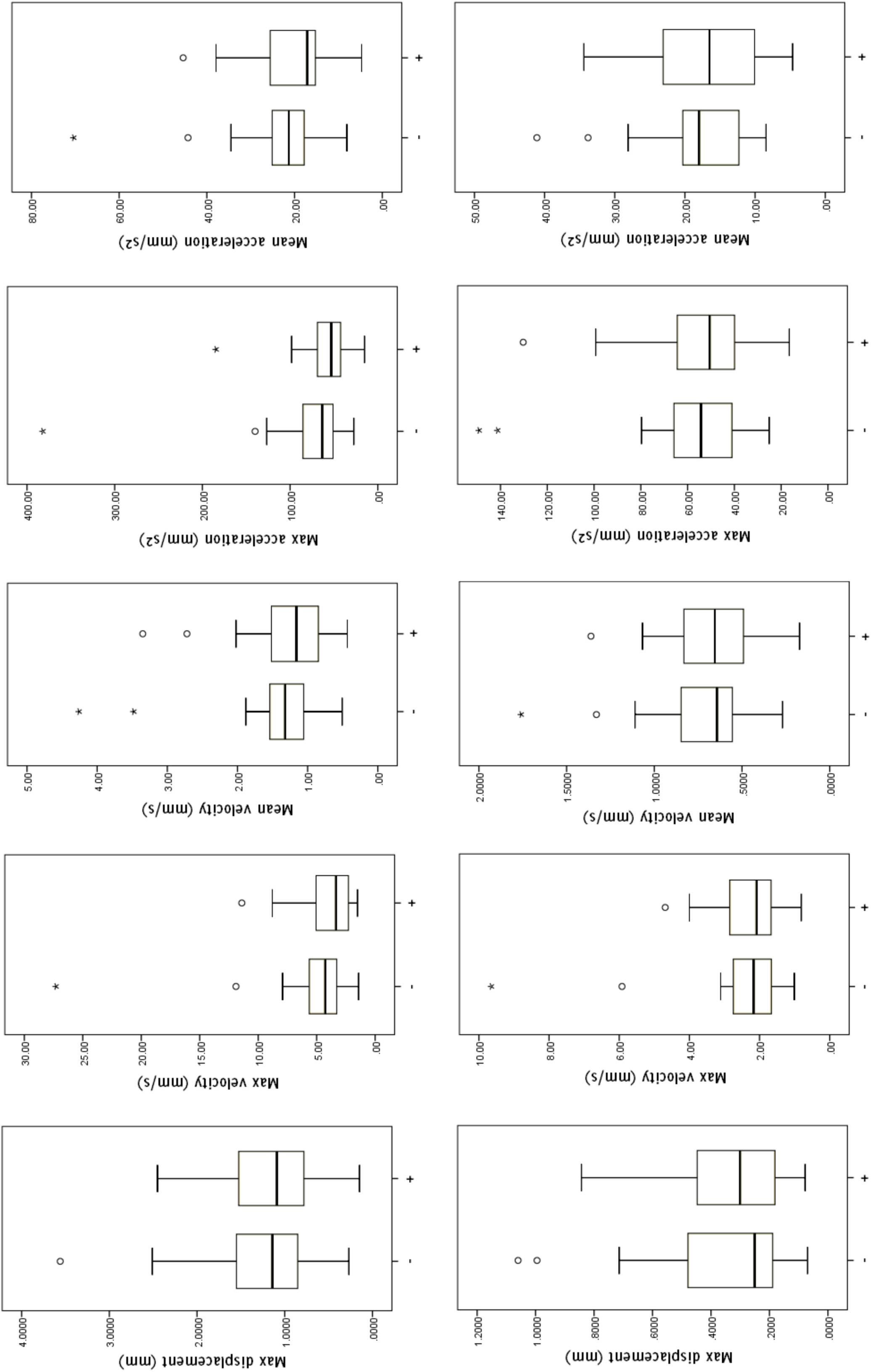
Box-whisker plots showing the distribution of the motion parameters (relative to the probe: top row, relative to the periadventitial tissues: bottom row) within the asymptomatic (marked -) and symptomatic (marked +) groups. Horizontal bars within boxes show median values.

**3 -.**
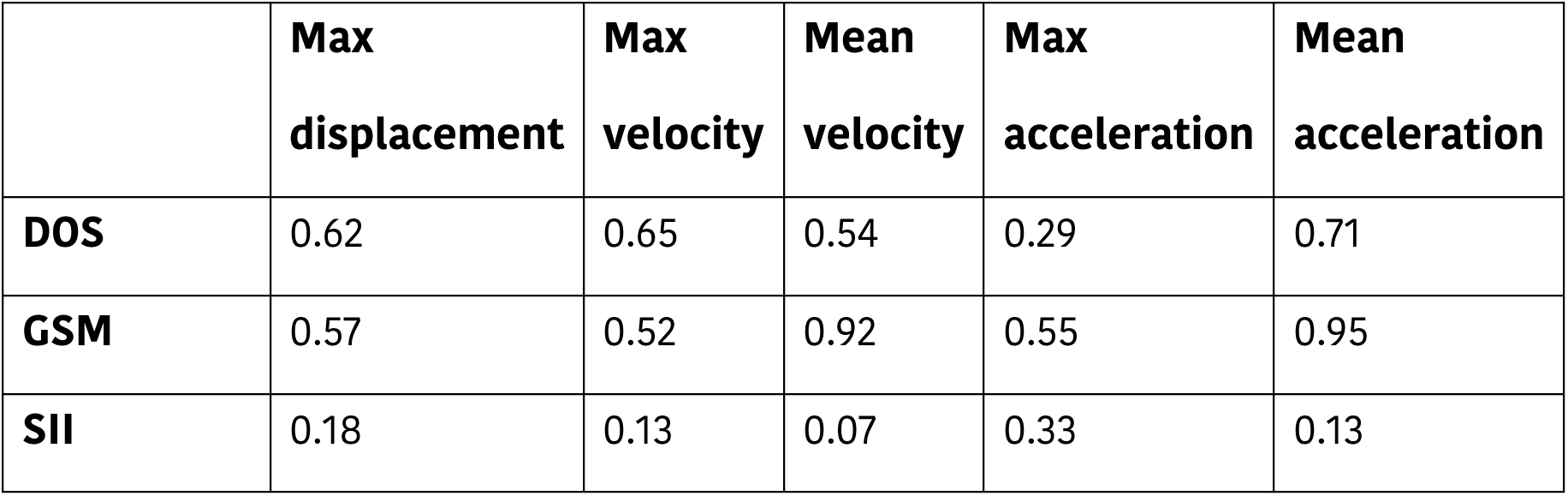
Significance of association (p-values) between motion parameters relative to the ultrasound probe, the degree of stenosis (DOS), plaque greyscale median (GSM) and the surface irregularity index (SII).

**4 -.**
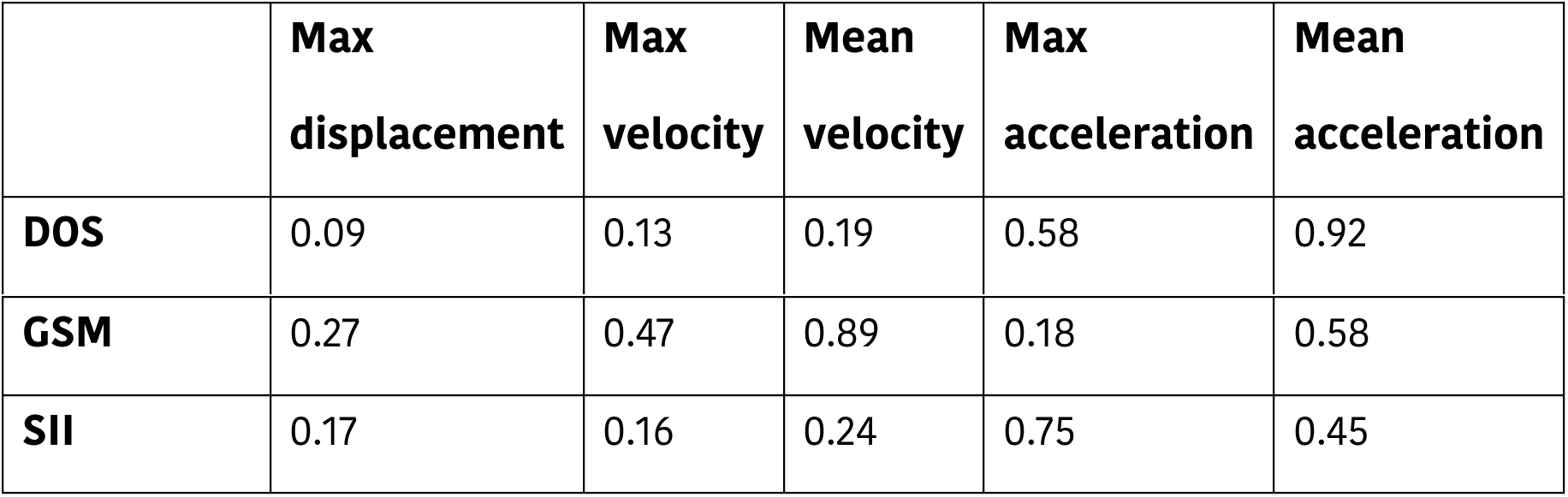
Significance of association (p-values) between motion parameters relative to the periadventitial tissues, the degree of stenosis (DOS), plaque greyscale median (GSM) and the surface irregularity index (SII).

|  | **Max displacement** | **Max velocity** | **Mean velocity** | **Max acceleration** | **Mean acceleration** |
| --- | --- | --- | --- | --- | --- |
| **DOS** | 0.09 | 0.13 | 0.19 | 0.58 | 0.92 |
| **GSM** | 0.27 | 0.47 | 0.89 | 0.18 | 0.58 |
| **SII** | 0.17 | 0.16 | 0.24 | 0.75 | 0.45 |

Measurement reproducibility was good for parameters representing the motion of the plaque relative to the probe (COV < 10%). It was better for mean plaque velocity and mean plaque acceleration (COV < 5%), than for maximum plaque displacement, maximum plaque velocity, and maximum plaque acceleration (COV >= 5%, Table 5). In the case of the motion of the plaque relative to the periadventitial tissues, reproducibility was lower for all motion parameters (COV > 15%). *In vitro* assessment showed that motion was not detected below a set displacement of 50 μm while measurement error was higher in the range 50 to 100 μm (mean error 37.0%), compared with that in the range 200 to 500 μm (mean error 5.4%, Table 6). Figures 5 and 6 show the calculated motion parameters for the case where the actuator was set to produce maximum displacements of 500 and 200 μm, respectively. Periodic, downward displacements of 476 and 188 μm are seen in Figures 5 and 6, respectively, with a small horizontal motion component in the case of Figure 6. In agreement with the programming of the actuator, velocity and acceleration were found to be greater when the motion was away from the ultrasound probe (increasing y values) compared to towards to probe (decreasing y values) (Figure 5).

**5 -.**
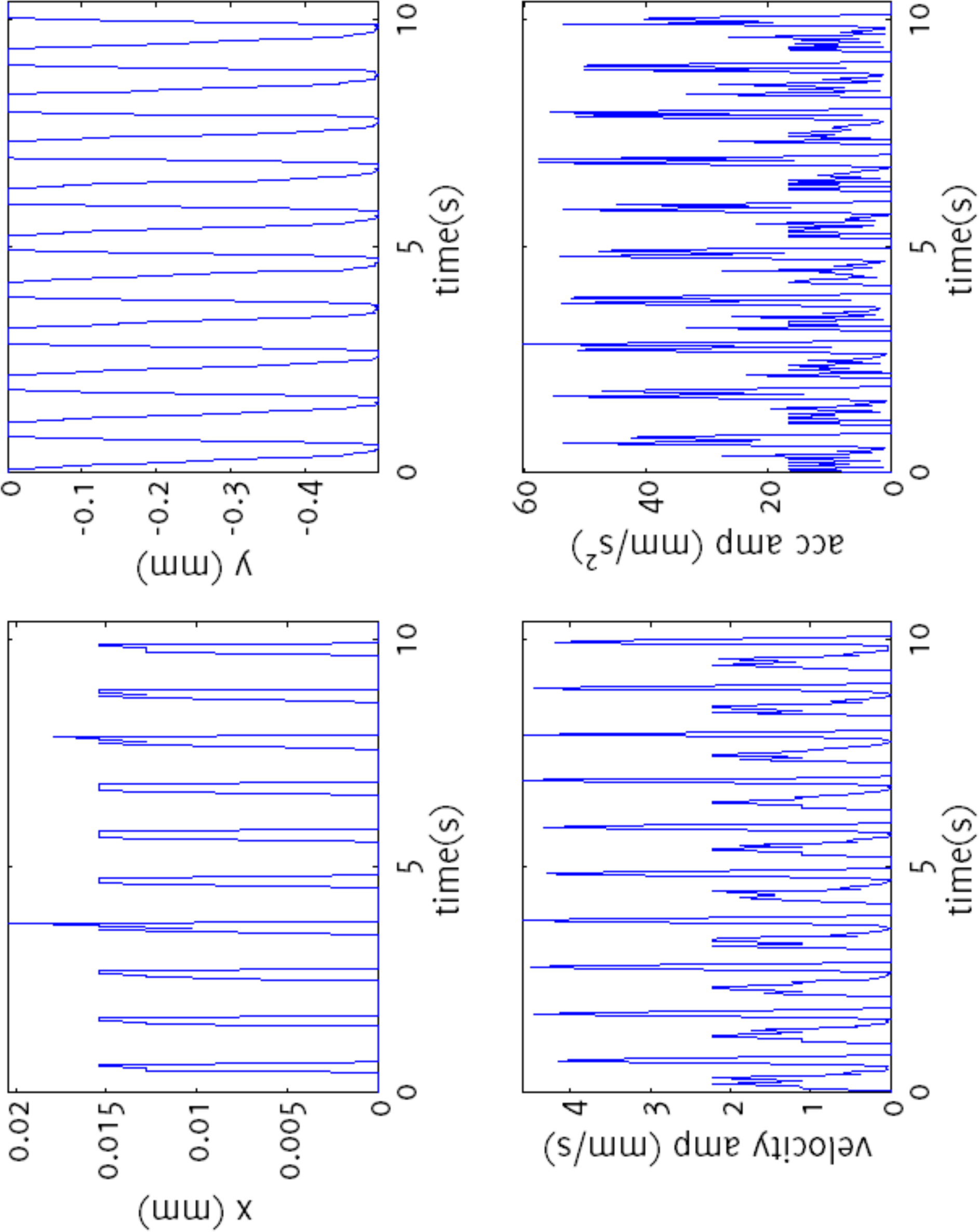
Calculated motion parameters for the *in vitro* study with the actuator set to produce a maximum displacement of 500 μm The horizontal and vertical components of position are shown in the top-left and top-right plots, respectively. Velocity and acceleration magnitudes are shown in the plots at the bottom. A small amount of displacement was detected in the horizontal direction even though set the motion was purely along the vertical axis.

**5 -.** Reproducibility of the motion parameters (intra-observer coefficients of variation in %).

|  |  |  |
| --- | --- | --- |
| **Relative to the ultrasound probe** | **Max displacement** | 7.6 |
|  | **Max velocity** | 7.0 |
|  | **Mean velocity** | 1.6 |
|  | **Max acceleration** | 4.9 |
|  | **Mean acceleration** | 1.1 |
| **Relative to the periadventitial tissues** | **Max displacement** | 20.4 |
|  | **Max velocity** | 16.4 |
|  | **Mean velocity** | 19.3 |
|  | **Max acceleration** | 16.0 |
|  | **Mean acceleration** | 21.1 |

**6 -.** *In vitro* assessment comparing the measured motion of the tissue mimicking material (TMM) with the set displacement of the actuator and the motion of the TMM-lumen interface measured using wall motion techniques [23].

|  |  |  |  |  |  |  |  |  |
| --- | --- | --- | --- | --- | --- | --- | --- | --- |
|  | **Set displacement ($\mu\text{m}$)** | 5 | 10 | 20 | 50 | 100 | 200 | 500 |
| **Measured motion of the TMM wall** | **Max displacement ($\mu\text{m}$)** | 7 | 15 | 26 | 53 | 105 | 201 | 503 |
|  | **Max velocity (mm/s)** | 0.16 | 0.14 | 0.27 | 0.55 | 0.60 | 1.17 | 3.99 |
|  | **Mean velocity (mm/s)** | 0.02 | 0.03 | 0.06 | 0.12 | 0.22 | 0.37 | 0.95 |
|  | **Max acceleration ($\text{mm/s}^2$)** | 3.3 | 2.4 | 4.2 | 8.0 | 8.1 | 14.7 | 47.4 |
|  | **Mean acceleration ($\text{mm/s}^2$)** | 0.50 | 0.76 | 1.25 | 2.0 | 2.3 | 3.5 | 11.0 |
| **Measured motion of TMM** | **Max displacement ($\mu\text{m}$) [error]** | 0<br>[100%] | 0<br>[100%] | 0<br>[100%] | 74<br>[48%] | 74<br>[26%] | 188<br>[6.0%] | 476<br>[4.8%] |
|  | **Max velocity** **(mm/s)** | 0 | 0 | 0 | 1.3 | 1.2 | 1.26 | 4.9 |
|  | **Mean velocity**<br>**(mm/s)** | 0 | 0 | 0 | 0.16 | 0.16 | 0.34 | 0.91 |
|  | **Max**<br>**acceleration**<br>**(mm/s<sup>2</sup>)** | 0 | 0 | 0 | 22.5 | 20.5 | 22.2 | 75.7 |
|  | **Mean**<br>**acceleration**<br>**(mm/s<sup>2</sup>)** | 0 | 0 | 0 | 5.2 | 5.0 | 6.1 | 13.0 |

**6 -.**
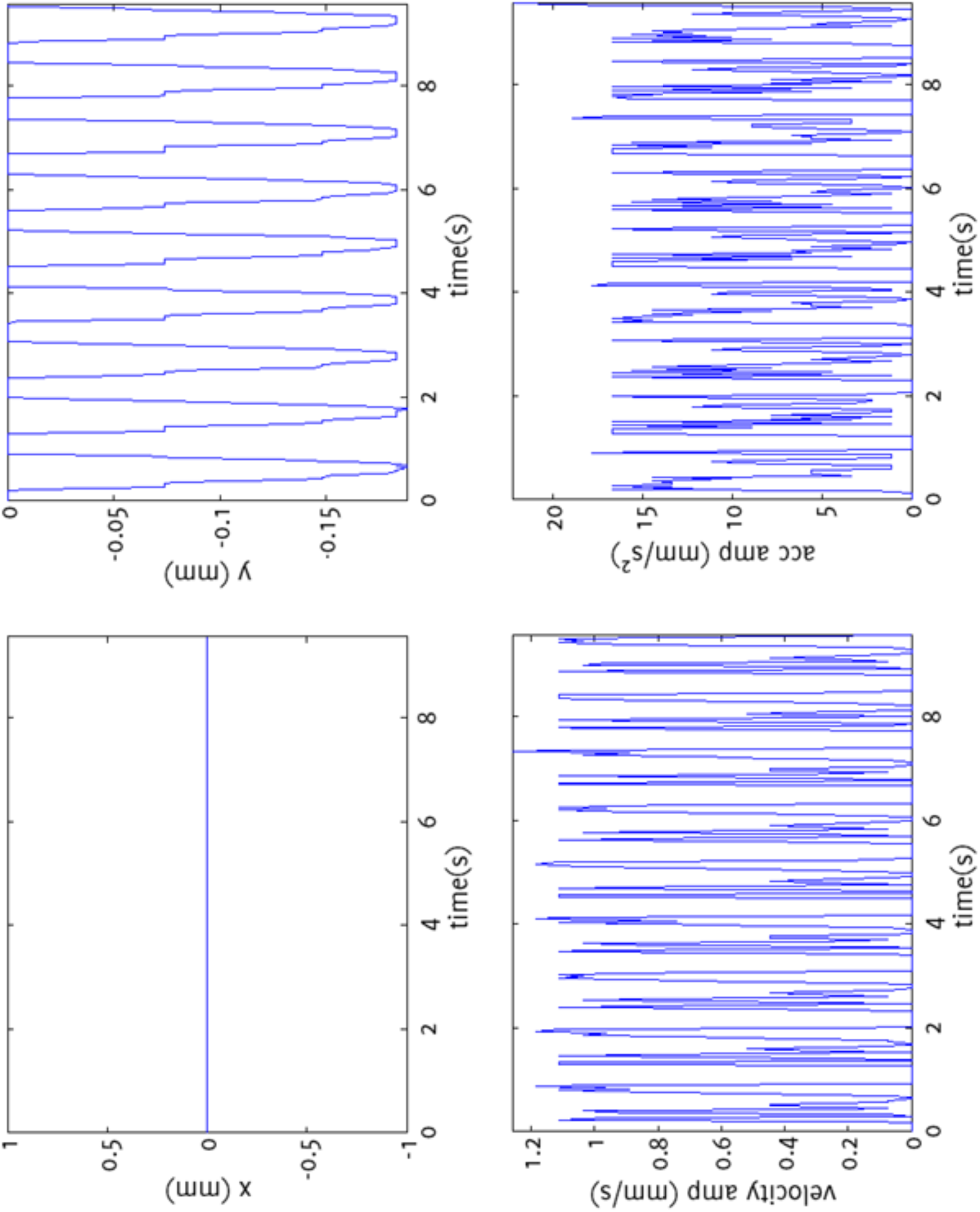
Calculated motion parameters for the *in vitro* study with the actuator set to produce a maximum displacement of 200 μm The horizontal and vertical components of position are shown in the top-left and top-right plots, respectively. Velocity and acceleration magnitudes are shown in the plots at the bottom.

## Discussion

This study provided an evaluation of the physiological motion of the carotid atherosclerotic plaque using B-Mode ultrasound image analysis, relative to the ultrasound probe and relative to the periadventitial tissues. Plaque velocities relative to the ultrasound probe have been investigated before, and our results are in accordance with previous findings [5,11]. Despite demonstrating the capability to detect motion amplitudes as low as 50 μm, we found no significant differences in any of the motion parameters in relation to the presence of cerebrovascular symptoms, and no significant association was found between plaque motion and the degree of carotid artery stenosis, or greyscale plaque characteristics. Nevertheless, this study provides useful data on physiological plaque motion in the carotid arteries.

Several studies have previously investigated the motion of the atherosclerotic plaque. Chan's early work confirmed the existence of carotid plaque motion when patient or probe motion is taken into account [44]. However, this was a preliminary study looking at only two clinical image sequences at low frame rates, and did not establish the presence of any discrepant motion between the plaque and the periadventitial tissues. Iannuzzi *et al*. used a qualitative assessment scheme based on an apparent distal shift of the plaque axis, and found that longitudinal plaque motion was associated with ipsilateral brain involvement in transient ischemic attack patients [1]. However, only a small percentage of the plaques (37%) were found to have longitudinal lesion motion in that study and the analysis was based on 18 plaques having longitudinal plaque motion in the artery ipsilateral to hemispheric damage, compared to 6 plaques which had longitudinal plaque motion in the artery contralateral to hemispheric damage. Other studies, on the other hand, reported that plaques from symptomatic patients had higher maximum discrepant surface velocities compared with plaques from asymptomatic patients [8,11].

Since no significant relationship was found between the parameters describing the motion of the plaque and the presence of cerebrovascular symptoms in this study, other complementary methods such as plaque characterisation [40-41,45], intraplaque strain analysis [26-28] and plaque elastography [46-49] may be suggested as avenues for research to help identify the vulnerable carotid plaque. However, this study had several limitations which may have hindered the unravelling of a possible association between plaque motion and vulnerability. The first was the relatively high rate of unsuccessful acquisitions, with speckle decorrelation accountable for 87% of the 15 unusable data sets. Large amounts of overlying tissue in obese patients was the main reason behind the speckle decorrelation in our acquisitions as the limited penetration depth of the ultrasound beam resulted in poor image quality. Ultrasonic shadowing, caused by calcified plaques, was the other aetiology of unsuccessful motion analyses. On the other hand, a success rate of 81% is similar to that of other studies investigating plaque motion, with similar underlying factors for unsuccessful analysis [11]. Secondly, although no association was found between plaque motion and the presence of cerebrovascular symptoms, it should be remembered that a comparison with plaque histology may have identified important associations as the presence of cerebrovascular symptoms is only a proxy for plaque vulnerability. Thirdly, since plaque motion is hypothesised to be a possible cause of plaque rupture, a prospective study looking at future incidence of cerebrovascular events can also identify important relationships. Therefore, plaque motion analysis, particularly the motion relative to the periadventitial tissues also remains an important research avenue for identifying the “at-risk” carotid atherosclerotic plaque.

## Conclusions

This study provided physiological data on carotid artery plaque bulk motion and motion relative to the periadventitial tissues. The assessment was performed using equipment readily available in the stroke clinic with a technique that was demonstrated to be capable of detecting motion amplitudes as small as 50μm. However, there were no significant differences found in the measured motion parameters with respect to the presence of cerebrovascular symptoms, the degree of stenosis, or the greyscale plaque characteristics.

## Competing interests

The authors declare that they have no competing interests.

## Authors' Contributions

The study was designed by KVR and BK. Patient recruitment was carried out by JWG. Ultrasound imaging data were collected by TCH and analysed by BK. All authors contributed to the interpretation and presentation of the results and, read and approved the final manuscript.

## Acknowledgements

The research was funded by and took place at the National Institute for Health Research (NIHR) Collaboration for Leadership in Applied Health Research and Care based at the University Hospitals of Leicester NHS Trust. The views expressed are those of the authors and not necessarily those of the NHS, the NIHR or the Department of Health.

## Additional Files

### Additional Video File 1

Plaque and periadventitial tissues with motion tracking. Arrows show the local displacement at time t with respect to the position at frame 1, magnified by a factor of 10. The individual motion vectors are averaged separately for the plaque body and the periadventitial tissues to provide sub-pixel resolution.

## References

1. Iannuzzi A, Wilcosky T, Mercuri M, Rubba P, Bryan FA, Bond MG. Ultrasonographic correlates of carotid atherosclerosis in transient ischemic attack and stroke. Stroke. 1995;26:614-619.

2. Woodcock JP. Characterisation of the atheromatous plaque in the carotid arteries. Clinical Physics and Physiological Measurement. 1989;10 Suppl A:45-49.

3. Falk E. Why do plaques rupture? Circulation. 1992;86:III30-III42.

4. Hennerici MG. The unstable plaque. Cerebrovascular Diseases. 2004;17 Suppl 3:17-22.

5. Bang J, Dahl T, Bruinsma A, Kaspersen JH, Nagelhus Hernes TA, Myhre HO. A new method for analysis of motion of carotid plaques from RF ultrasound images. Ultrasound in Medicine & Biology. 2003;29:967-976.

6. Ogata T, Yasaka M, Wakugawa Y, Kitazono T, Okada Y. Morphological classification of mobile plaques and their association with early recurrence of stroke. Cerebrovascular Diseases. 2011;30:606-611.

7. Kume S, Hama S, Yamane K, Wada S, Nishida T, Kurisu K. Vulnerable carotid arterial plaque causing repeated ischemic stroke can be detected with B-mode ultrasonography as a mobile component: Jellyfish sign. Neurosurgical Review. 2011;33:419-430.

8. Stoitsis J, Golemati S, Nikita KS, Nicolaides AN. Characterization of carotid atherosclerosis based on motion and texture features and clustering using fuzzy c-means. Proceedings of the IEEE Engineering in Medicine and Biology Society. 2007;2:1407-1410.

9. Nasrabadi H, Pattichis MS, Fisher P, Nicolaides AN, Griffin M, Makris GC, et al. Measurement of motion of carotid bifurcation plaques. Proceedings of the IEEE International Conference on Bioinformatics & Bioengineering. 2012.

10. Kashiwazaki D, Yoshimoto T, Mikami T, Muraki M, Fujimoto S, Abiko K, et al. Identification of high-risk carotid artery stenosis: motion of intraplaque contents detected using B-mode ultrasonography. Journal of Neurosurgery. 2013;117:574-578.

11. Meairs S, Hennerici M. Four-dimensional ultrasonographic characterization of plaque surface motion in patients with symptomatic and asymptomatic carotid artery stenosis. Stroke. 1999;30:1807-1813.

12. Svedlund S, Gan L. Longitudinal wall motion of the common carotid artery can be assessed by velocity vector imaging. Clinical Physiology and Functional Imaging. 2011;31:32-38.

13. Bukaeraction computational model, and experimental validation. Mathematical Biosciences and Engineering. 2013;10:295-318.

14. Zahnd G, Orkisz M, Sérusclat A, Moulin P, Vray D. Evaluation of a Kalman-based block matching method to assess the bi-dimensional motion of the carotid artery wall in B-mode ultrasound sequences. Medical Image Analysis. 2014;17:573-585.

15. Zahnd G, Vray D, Sérusclat A, Alibay D, Bartold M, Brown A, et al. Longitudinal displacement of the carotid wall and cardiovascular risk factors: associations with aging, adiposity, blood pressure and periodontal disease independent of crosssectional distensibility and intima-media thickness. Ultrasound in Medicine & Biology. 2013;38:1705-1715.

16. Ahlgren AR, Cinthio M, Persson HW, Lindström K. Different patterns of longitudinal displacement of the common carotid artery wall in healthy humans are stable over a four-month period. Ultrasound in Medicine & Biology. 2012;38:916-925.

17. Ahlgren AR, Cinthio M, Steen S, Nilsson T, Sjöberg T, Persson HW, et al. Longitudinal displacement and intramural shear strain of the porcine carotid artery undergo profound changes in response to catecholamines. American Journal of Physiology. Heart and Circulatory Physiology. 2012;302:H1102-H1115.

18. Cinthio M, Ahlgren AR. Intra-observer variability of longitudinal displacement and intramural shear strain measurements of the arterial wall using ultrasound noninvasively in vivo. Ultrasound in Medicine & Biology. 2010;36:697-704.

19. Ahlgren AR, Cinthio M, Steen S, Persson HW, Sjöberg T, Lindström K. Effects of adrenaline on longitudinal arterial wall movements and resulting intramural shear strain: a first report. Clinical Physiology and Functional Imaging. 2009;29:353-359.

20. Cinthio M, Ahlgren AR, Bergkvist J, Jansson T, Persson HW, Lindström K. Longitudinal movements and resulting shear strain of the arterial wall. American Journal of Physiology. Heart and Circulatory Physiology. 2006;291:H394-H402.

21. Cinthio M, Ahlgren AR, Jansson T, Eriksson A, Persson HW, Lindström K. Evaluation of an ultrasonic echo-tracking method for measurements of arterial wall movements in two dimensions. IEEE Transactions on Ultrasonics Ferroelectrics and Frequency Control. 2005;52:1300-1311.

22. Persson M, Ahlgren AR, Jansson T, Eriksson A, Persson HW, Lindström K. A new non-invasive ultrasonic method for simultaneous measurements of longitudinal and radial arterial wall movements: first in vivo trial. Clinical Physiology and Functional Imaging. 2003;23:247-251.

23. Kanber B, Hartshorne TC, Horsfield MA, Naylor AR, Robinson TG, Ramnarine KV. Wall motion in the stenotic carotid artery: association with greyscale plaque characteristics, the degree of stenosis and cerebrovascular symptoms. Cardiovascular Ultrasound. 2013;11:37.

24. Golemati S, Sassano A, Lever MJ, Bharath AA, Dhanjil S, Nicolaides AN. Carotid artery wall motion estimated from B-mode ultrasound using region tracking and block matching. Ultrasound in Medicine & Biology. 2003;29:387-399.

25. Stoitsis J, Golemati S, Dimopoulos A, Nikita K. Analysis and quantification of arterial wall motion from B-mode ultrasound images - comparison of block-matching and optical flow. Conf Proc IEEE Eng Med Biol Soc. 2007;5:4469-4472.

26. McCormick M, Varghese T, Wang X, Mitchell C, Kliewer MA, Dempsey RJ. Methods for robust in vivo strain estimation in the carotid artery. Physics in Medicine and Biology. 2012;57:7329-7353.

27. Shi H, Mitchell CC, McCormick M, Kliewer MA, Dempsey RJ, Varghese T. Preliminary in vivo atherosclerotic carotid plaque characterization using the accumulated axial strain and relative lateral shift strain indices. Physics in Medicine and Biology. 2008;53:6377-6394.

28. Rocque BG, Jackson D, Varghese T, Hermann B, McCormick M, Kliewer M, et al. Impaired cognitive function in patients with atherosclerotic carotid stenosis and correlation with ultrasound strain measurements. Journal of the Neurological Sciences. 2013;322:20-24.

29. Naim C, Cloutier G, Mercure E, Destrempes F, Qin Z, El-abyad W, et al. Characterisation of carotid plaques with ultrasound elastography: feasibility and correlation with high-resolution magnetic resonance imaging. European Radiology. 2014;23:2030-2041.

30. Schmitt C, Soulez G, Maurice RL, Giroux M, Cloutier G. Noninvasive vascular elastography: toward a complementary characterization tool of atherosclerosis in carotid arteries. Ultrasound in Medicine & Biology. 2007;33:1841-1858.

31. De Korte CL, Hansen HHG, Van Der Steen AFW. Vascular ultrasound for atherosclerosis imaging. Interface Focus. 2012;1:565-575.

32. Ribbers H, Lopata RGP, Holewijn S, Pasterkamp G, Blankensteijn JD, De Korte CL. Noninvasive two-dimensional strain imaging of arteries: validation in phantoms and preliminary experience in carotid arteries in vivo. Ultrasound in Medicine & Biology. 2007;33:530-540.

33. Hasegawa H, Kanai H, Ichiki M, Tezuka F. Ultrasonic measurement of arterial wall for quantitative diagnosis of atherosclerosis‐‐elasticity imaging of arterial wall and atherosclerotic plaque. The Japanese Journal of Clinical Pathology. 2007;55:363-368.

34. Lenzi GL, Vicenzini E. The ruler is dead: an analysis of carotid plaque motion. Cerebrovascular Diseases. 2006;23:121-125.

35. Teirlinck CJ, Bezemer RA, Kollmann C, Lubbers J, Hoskins PR, Ramnarine KV, et al. Development of an example flow test object and comparison of five of these test objects, constructed in various laboratories. Ultrasonics. 1998;36:653-660.

36. Ramnarine KV, Anderson T, Hoskins PR. Construction and geometric stability of physiological flow rate wall-less stenosis phantoms. Ultrasound in Medicine & Biology. 2001;27:245-250.

37. North American Symptomatic Carotid Endarterectomy Trial Collaborators. Beneficial effect of carotid endarterectomy in symptomatic patients with high-grade carotid stenosis. The New England Journal of Medicine. 1991;325:445-453.

38. Grant EG, Benson CB, Moneta GL, Alexandrov AV, Baker JD, Bluth EI, et al. Carotid artery stenosis: gray-scale and Doppler US diagnosis‐‐Society of Radiologists in Ultrasound Consensus Conference. Radiology. 2003;229:340-346.

39. Oates CP, Naylor AR, Hartshorne T, Charles SM, Fail T, Humphries K, et al. Joint recommendations for reporting carotid ultrasound investigations in the United Kingdom. European Journal of Vascular and Endovascular Surgery. 2008;37:251-261.

40. Kanber B, Hartshorne TC, Horsfield MA, Naylor AR, Robinson TG, Ramnarine KV. Dynamic variations in the ultrasound greyscale median of carotid artery plaques. Cardiovascular Ultrasound. 2013;11:21.

41. Kanber B, Hartshorne TC, Horsfield MA, Naylor AR, Robinson TG, Ramnarine KV. Quantitative assessment of carotid plaque surface irregularities and correlation to cerebrovascular symptoms. Cardiovascular Ultrasound. 2013;11:38.

42. Kanber B, Ramnarine KV. A Probabilistic Approach to Computerized Tracking of Arterial Walls in Ultrasound Image Sequences. ISRN Signal Processing. 2012.

43. Ramnarine KV, Kanber B, Panerai RB. Assessing the performance of vessel wall tracking algorithms: the importance of the test phantom. Journal of Physics Conference Series. 2004;1:199-204.

44. Chan KL. Two approaches to motion analysis of the ultrasound image sequence of carotid atheromatous plaque. Ultrasonics. 1993;31:117-123.

45. Kanber B, Hartshorne TC, Horsfield MA, Naylor AR, Robinson TG, Ramnarine KV. A Novel Ultrasound-Based Carotid Plaque Risk Index Associated with the Presence of Cerebrovascular Symptoms. Ultraschall in der Medizin. 2014.

46. Ramnarine KV, Garrard JW, Dexter K, Nduwayo S, Panerai RB, Robinson TG. Shear wave elastography assessment of carotid plaque stiffness: in vitro reproducibility study. Ultrasound in Medicine & Biology. 2013;40:200-209.

47. Garrard JW, Ramnarine KV. Shear-Wave Elastography in Carotid Plaques: Comparison with Grayscale Median and Histological Assessment in an Interesting Case. Ultraschall in der Medizin. 2013.

48. Ramnarine KV, Garrard JW, Kanber B, Nduwayo S, Hartshorne TC, Robinson TG. Shear wave elastography imaging of carotid plaques: feasible, reproducible and of clinical potential. Cardiovascular Ultrasound. 2014;12:49.

49. Garrard JW, Ummur P, Nduwayo S, Kanber B, Hartshorne TC, West KP, et al. Shear Wave Elastography May Be Superior to Greyscale Median for the Identification of Carotid Plaque Vulnerability: A Comparison with Histology. Ultraschall in der Medizin. 2015;36:386-390.

